# Deep generative networks reveal the tuning of neurons in IT and predict their influence on visual perception

**DOI:** 10.1101/2024.10.09.617382

**Authors:** Paolo Papale, Daniela De Luca, Pieter R. Roelfsema

**Affiliations:** Department of Vision & Cognition, Netherlands Institute for Neuroscience (KNAW), 1105 BA Amsterdam, Netherlands; The BioRobotics Insistute and Department of Excellence in Robotics and AI, Scuola Superiore Sant’Anna, Pisa, Italy; Department of Integrative Neurophysiology, VU University, De Boelelaan 1085, 1081 HV Amsterdam, Netherlands; Department of Neurosurgery, Academic Medical Centre, Postbus 22660, 1100 DD Amsterdam, Netherlands; Laboratory of Visual Brain Therapy, Sorbonne Université, INSERM, CNRS, Institut de la Vision, 17 rue Moreau, F-75012 Paris, France

## Abstract

Finding the tuning of visual neurons has kept neuroscientists busy for decades. One approach to this problem has been to test specific hypotheses on the relevance of a visual property (e.g. orientation or color), build a set of “artificial” stimuli that vary along that property and then record neural responses to those stimuli. Here, we present a complementary, data-driven method to retrieve the tuning properties of visual neurons. Exploiting deep generative networks and electrophysiology in monkeys, we first used a method to reconstruct any stimulus from its evoked neuronal activity in the inferotemporal cortex (IT). Then, by arbitrarily perturbing the response of individual cortical sites in the model, we generated naturalistic and interpretable sequences of images that strongly influence neural activity of that site. This method enables the discovery of previously unknown tuning properties of high-level visual neurons that are easily interpretable, which we tested with carefully controlled stimuli. When we knew which images drove the neurons, we activated the cells with electrical microstimulation and observed a predicable shift of the monkey perception in the direction of the preferred image. By allowing the brain to tell us what it cares about, we are no longer limited by our experimental imagination.

---

Although it feels effortless to us, interpreting a daily visual scene is a complex and computationally intensive task for our brain. Consider the cloud of dots in Fig.1A, representing the set of all possible images we may face in a life-time. Each point in this cloud represents an image: one random point could portray a giraffe, another a toucan or a landscape, etc. More similar images are closer to each other, and dissimilar images are farther apart. This cloud is theoretically infinite, but despite the huge size of this set, we perceive completely new, unseen stimuli with very little ambiguity. How can our brain represent this large set of images with a relatively small number of neurons? One hypothesis is that the brain maps natural images into a more manageable, low-dimensional representation: a coordinate system. We can visualize a single dimension of this coordinate system as an axis, spanning the visual set from side to side. The direction of the axis tells us how much of a certain property is present in each image. For example, in Fig.1B, different images of the same building under different lighting conditions are arranged along the axis depending on their level of luminance/brightness.

The activity of visual neurons depends on the presence and magnitude of a specific property in the image that they are “tuned” to. Therefore, the activity of some neurons in a region of the brain sensitive to luminance would be high in response to the brightest image, and lower in response to the darkest image (Fig.1C), but other neurons might have the opposite preference. With a few axes that correspond to different visual properties, the brain could represent many images, each with a different coordinate in the representational space (Fig.1D).

Finding the tuning of visual neurons has kept neuroscientists busy for decades: this process requires to link the observed responses of visual neurons to some specific physical property of the stimulus^1^. Traditionally, this problem has been approached using a theory driven perspective: spending a lot of time on the blackboard, thinking of a possible property of interest, then build a set of “artificial” stimuli that vary along that property (e.g. orientation or color) and then record neural responses to those stimuli. Fig. 1E shows examples of controlled stimuli linearly varying along orientation, contrast or object curvature^2^. Studies employing artificial stimuli have provided important insight into neuronal and represent a critical test of our knowledge on the visual system.

**Figure 1.**
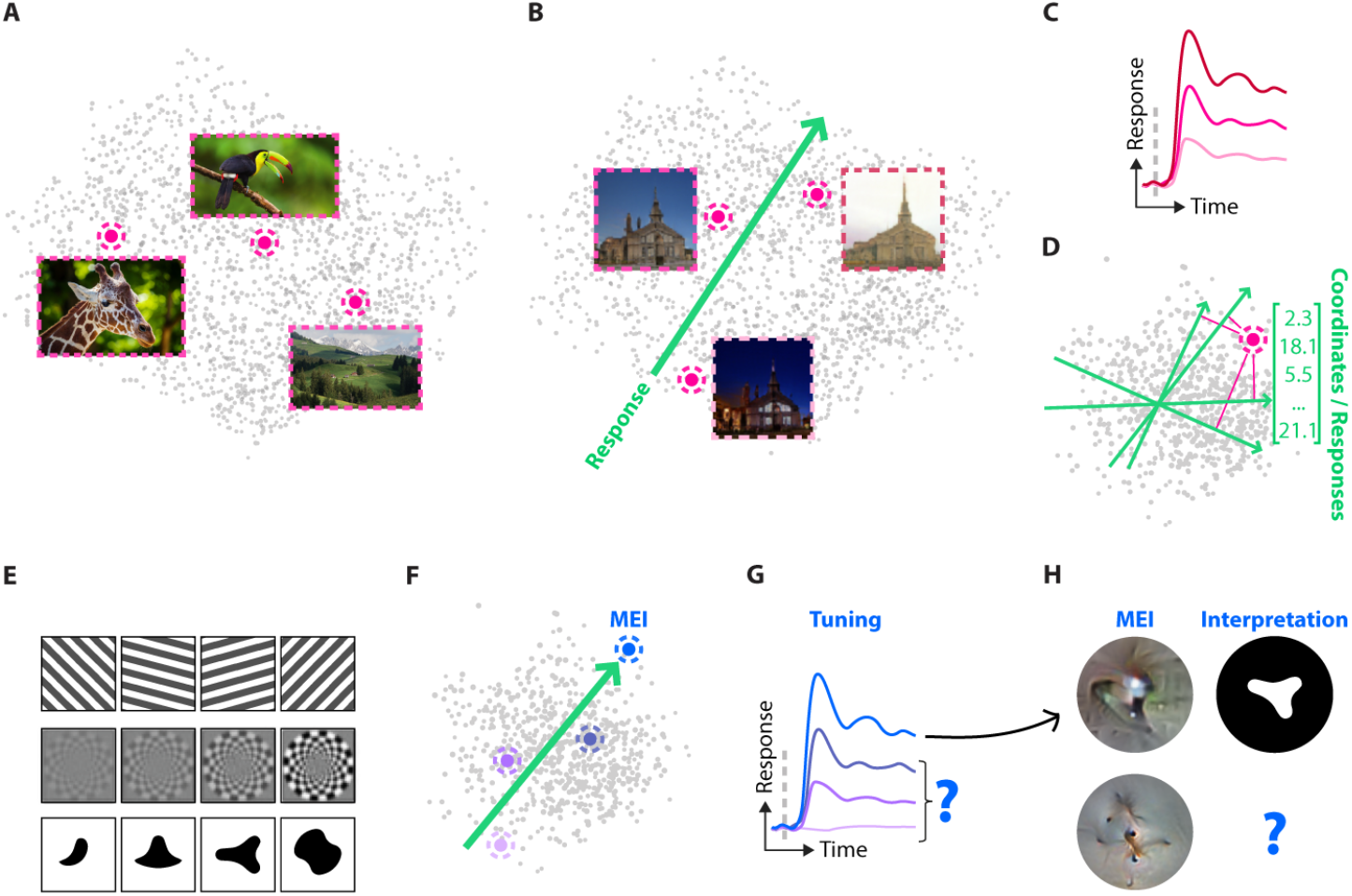
Linking visual properties to the tuning of neurons. Visual processing relies on intensive computations and a large portion of our brain, thus, understanding how vision works has kept neuroscientists busy for decades. **(A)** Each point in this cloud represents every possible visual input that we might encounter in our life. **(B)** Dots are arranged along axis representing a single visual property, e.g. brightness. **(C)** Visual neurons are tuned to specific visual properties and vary their response depending on its magnitude. **(D)** By relying on neurons tuned to different properties, the brain can represent every possible dot of the cloud with extreme reliability. **(E)** Neuroscientists have studied such tuning properties using carefully designed artificial stimuli. **(F)** A most-exciting image (MEI), produced by a data-driven method, can be represented as the dot at the end of the response vector of a specific neuron. **(G)** MEIs correspond to the maximum response (light blue line), but they do not capture the tuning of the neuron. **(H)** The interpretability of MEIs is difficult. It is unknown whether the top MEI is related to an image property like curvature, and it might be even more difficult to derive the image property that is related to the bottom MEI.

However, as scientists facing such a complex problem as understanding the brain, we are challenged by a simple, at times hurting truth: we are limited by our experimental imagination. For this reason, a different approach that complements the theory driven paradigm has recently gained popularity, thanks to the availability of more powerful computing resources. This second approach is data driven, as it does not rely on explicit theories and carefully designed stimuli, but instead the link between the physical properties of the visual input and the tuning of visual neurons is inferred directly from their neuronal responses. Recent data-driven approaches relied on recording neural activity in response to ecological stimuli, and then detecting meaningful patterns using computational models^3,4^. These models learn a mapping between the activations of an artificial neural network (ANN) and the responses of neurons to thousands of images, that can be used to produce images that maximize the response of individual neurons^5–7^. When these so-called most exciting inputs (MEIs) are shown to neurons they can drive the activity even beyond the usual range^5–7^ (Figure 1F).

Using ANNs to understand tuning has proven a powerful tool to study the highly non-linear tuning properties in high-level regions such as area V4 and the inferotemporal cortex (IT)^5,7,8^. However, this data-driven method also has some limitations. First, this method can produce a single image that elicits strong activity of neurons, but it is not aimed at revealing the full tuning of a neuron (with the exception of ref.^9^; Figure 1F-G). Second, many MEIs are hard to interpret because they do not reveal how much of the neuron’s response depend on the various features of the MEI (Figure 1H).

An ideal data-driven method to discover new tuning properties should produce interpretable descriptions of the tuning, like image sequences that cause the largest change in the response. These interpretable sequences should be easy to translate into image-computable metrics, that can predict the response of neurons, groups of neurons or voxels in fMRI. We present here a proof-of-concept approach that achieves such a goal. We also demonstrate that this data-driven approach predicts the effects of microstimulation on visual perception.

## Data-driven measurement of the tuning of IT neurons

The approach that we present here is a simple extension of a state-of-the-art decoding method for reconstructing seen images from their evoked neural responses^10^. We exploited a Generative Adversarial Network (GAN), which was trained to generate natural-looking images from “latent vectors”, which are compressed representation of visual scenes. The latent vectors are defined by their coordinates in a latent space, akin to the cloud of dots described in Figure 1. A GAN is composed by a Generator (Figure 2A, blue) and Discriminator network (Figure 2A, pink) that are trained adversarially. The goal of the Discriminator is to differentiate between real images (Figure 2A, grey) and fake images produced by the Generator and the goal of the Generator is to produce images that look natural enough to fool the Discriminator. Improvements in one network pushes the other one to also improve.

**Figure 2.**
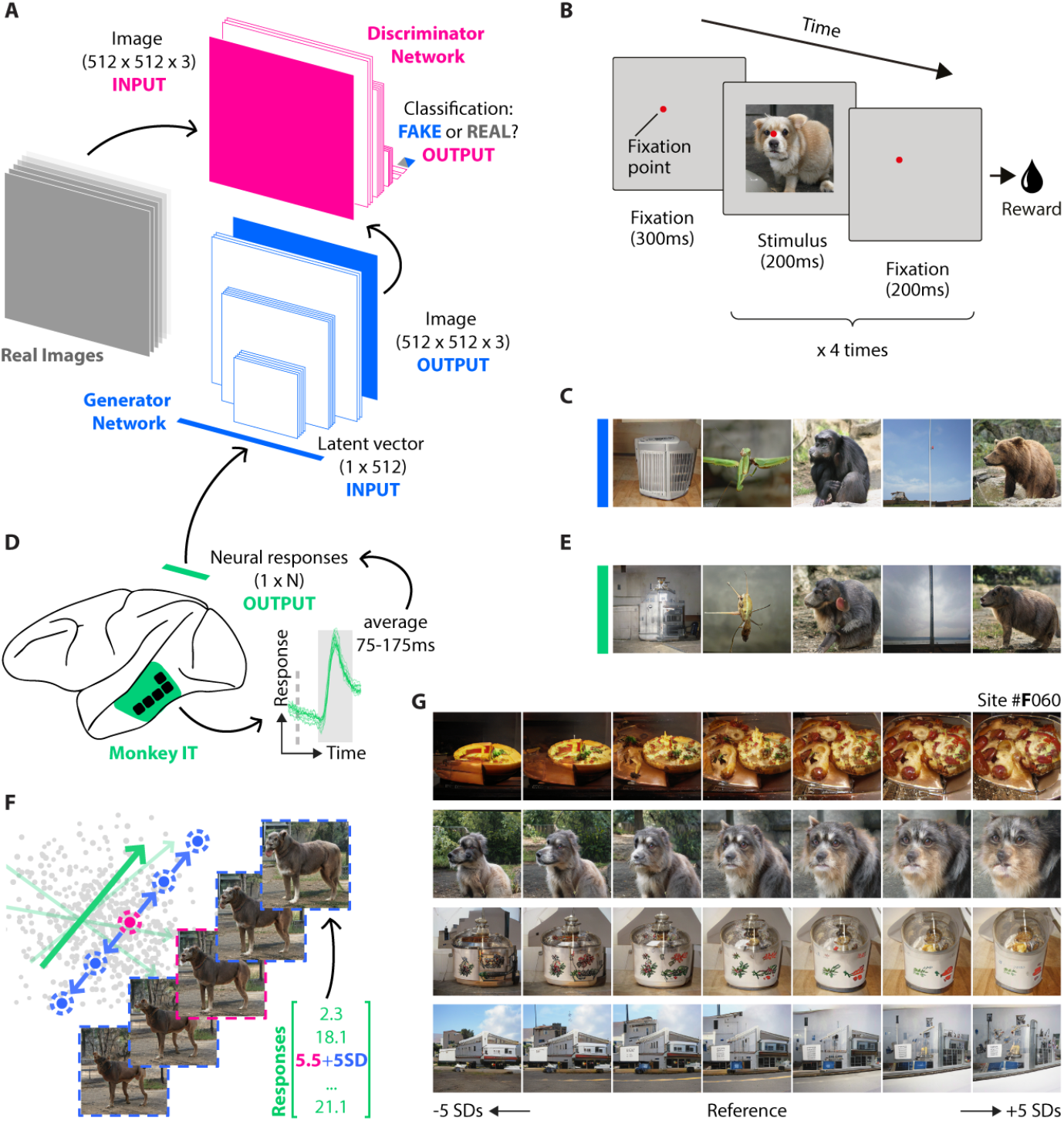
Data-driven measurement of the tuning of IT neurons. We used a Generative Adversarial Network (GAN) to create stimuli that were presented to two monkeys. We then used the GAN to study the relation between neuronal activity and the reconstructed stimulus. **(A)** A GAN is composed of two networks, a Generator and a Discriminator. The Generator produces images starting from a latent vector, which represents coordinates in a latent space (c.f. the dots in Figure 1A). **(B)** We presented the GAN to create stimuli that we showed for 200ms (followed by 200ms grey) to 2 monkeys fixating a red dot at the centre of the screen. **(C)** Five examples of GAN-generated stimuli. **(D)** We recorded the neural responses to the GAN-generated stimuli using chronically implanted electrodes in the inferotemporal cortex (IT). We determined a linear mapping between the pattern of neuronal responses (i.e. the neuronal vector) and the latent vector. This linear relationship allowed us to reconstruct the stimuli seen by the monkey (but left out from the training phase) from the activity in IT. **(E)** Stimulus reconstruction of the stimuli in **(C). (F)** The linear mapping allows us also to study the relation between MUA at individual recording sites and the reconstructed stimuli. By adding or subtracting activity (up to 5 SDs of each site’s response) we created images that illustrate the putative tuning of the neurons (tuning dimensions). **(G)** Example stimulus perturbations for a representative IT site, based on 4 reference images.

Every latent vector of a GAN corresponds to a unique image. The ability to generate images from latent vectors is helpful when the aim is to decode images that are presented to a subject from the neural responses elicited by them. In the present study we used a GAN with a latent vector with 512 elements to generate images with 512*512 pixels. We sampled a total of 4200 latent vectors, generated the associated images and showed them to two monkeys (monkey N and monkey F).

The monkeys fixated a dot at the center of the screen. After a delay of 300 ms, four images were presented for 200ms each, and they were all followed a grey screen that was visible for 200ms (Figure 2B). A few example images are shown in Figure 2C. We recorded the response elicited by these images with chronically implanted intracortical electrodes (Utah arrays) in posterior IT (Figure 2D). We recorded multi-unit activity (MUA) from 75 recordings sites in monkey N and 161 sites in monkey F, in response to 8,000 images shown in 2 recording sessions to each monkey.

We created vectors of the neuronal responses by combining the responses across all IT recording sites to each of the images in each of the monkeys, averaging the MUA in a time window from 75 to 275ms after stimulus onset. We included this vector of neural responses and the latent vector in a regression analysis to derive a linear mapping between the neuronal responses and the latent vectors. We used the response to 4000 images (shown once) for the linear regression, and used the response to 200 images that had been presented 20 times for the cross-validated generation of the images that were held out during training. Figure 2E shows that the reconstructions resembled the original images of Figure 2C. They had the same sematic class, and the shapes and colors were also similar. Accordingly, the correlation between the predicted latent vectors and the original ones was high in both monkeys (monkey N: Pearson’s r = 0.68±0.09; mean ± SD across recording sites and images, N = 75 sites, p < 0.001, t-test; monkey F: r = 0.73±0.08, N = 161 sites, p < 0.001, t-test; as in ref.^10^).

These results are in accordance with our previous findings^10^ that a vector of neural responses in IT can be used to reliably reconstruct the images that elicited these responses. We next examined how the images changed if we perturbed one element of the neural vector, varying the response of an individual cortical site while keeping the activity of the neurons at the other sites constant. We created sequences of 7 images by linearly adding or subtracting up to 5 standard deviations (SDs) of activity for each site, computed using the response to the 4000 train images (Figure 2F). The image perturbations provide insight into the stimulus dimensions that influence the neuronal activity at a particular recording site, and it is of interest to examine whether these perturbations are consistent across reference images. Figure 2G illustrates the image perturbations of 4 reference images for an example site (#F060). The model suggested that the activity of the neurons at the recording site increase when the central region of the image is zoomed in, which may be related to the spatial frequency content of this region.

## GAN generated image sequences reveal the tuning of IT neurons

The sequences of images that were generated by the GAN reflected known and previously reported tuning properties of IT neurons. A few representative examples are shown in Figure 3A. From top to bottom, the neurons appeared to be tuned to 1) object roundness, in line with previous reports showing tuning to contour curvature and shape complexity^11–13^, 2) colour saturation or contrast^14^, 3) the object perceived (or real-world) size^15^ 4) or its retinal size^16^. So far, the labelling of the sequences in Figure 3A was still speculative. Also, these seemed to change also according to other properties, even if less clearly, e.g. shape also appeared to change in the second sequence from the top. Thus, we needed further experimental testing to prove that this method is a useful source for advancing our knowledge on tuning properties.

**Figure 3.**
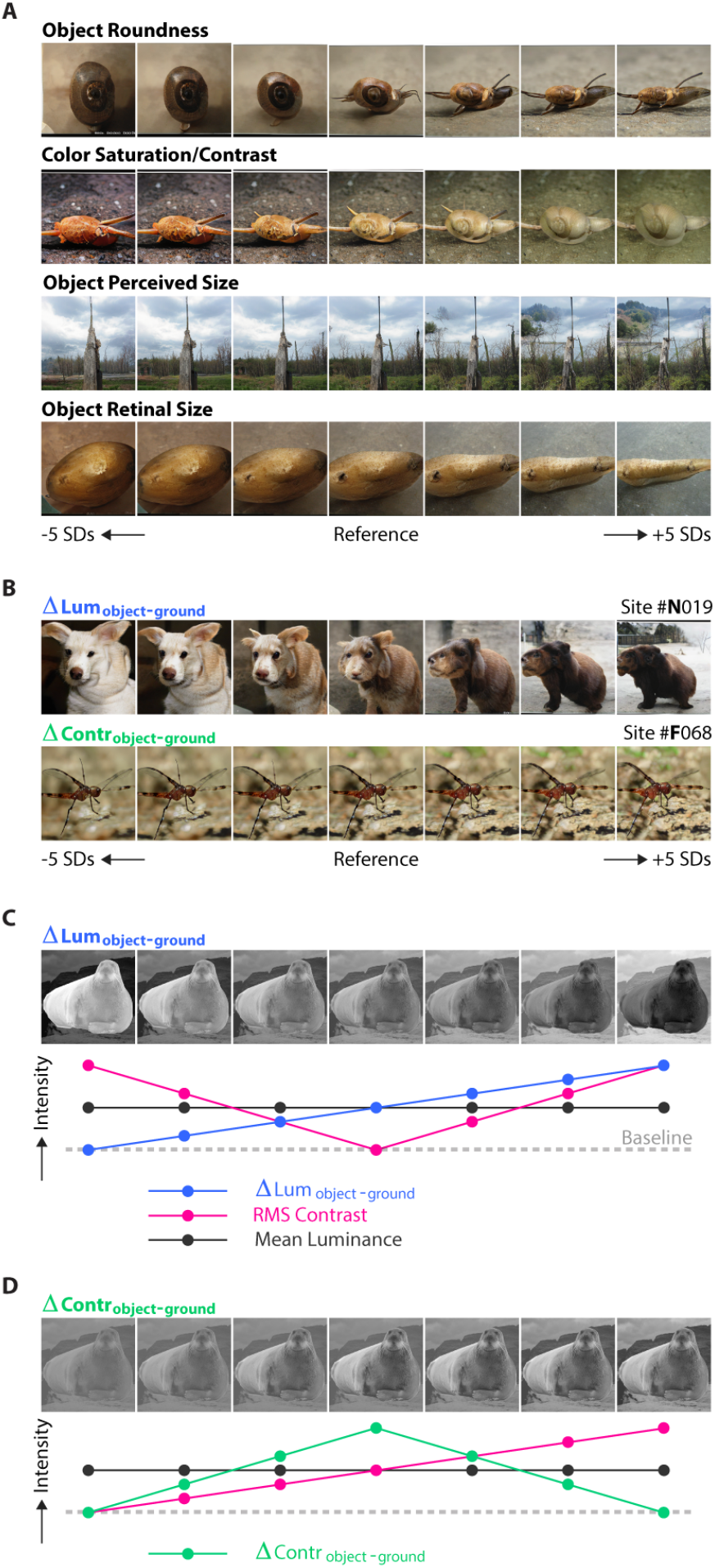
Feature dimensions that influence IT neurons and controlled stimuli for testing their influence. The GAN model reveals putative feature dimensions that influence the activity of IT neurons. **(A)** GAN-generated image sequences reveal image properties to which IT neurons are tuned. From top to bottom: object roundness, color saturation/contrast, object perceived (or real-world) size and object size. **(B)** Image sequences suggesting tuning of IT neurons to the relative luminance of the foreground object and the background (upper row) or the contrast of the background. Inspired by these sequences of images we built carefully controlled sequences of 7 stimuli. **(C)** Controlled stimuli in which we varied the luminance of the foreground object and the background in opposite directions. Lower panel: the intensity of ΔLum_object-ground_ (blue), global luminance (black) and RMS Contrast (pink) that we used as a reference known property and that was orthogonal to the novel dimension. **(D)** same as **(C)** for ΔContr_object-ground_.

We also observed IT neurons which appeared to be tuned to features, related to contrast and luminance of the foreground and background objects in the picture (Figure 3B). For example, the activity of neurons at site #N019 (Figure 3B, top) was, according to the GAN model, related to the relative luminance of the object and the background, i.e. the contrast polarity. The model predicted that the neurons would be most active for objects with a low luminance combined with backgrounds with a high luminance. Another example is given by recording site #F068 (Figure 3B, bottom). The model predicted that the neurons were most strongly driven if the background contrast is high.

In the examples of Figure 3B, there are more differences between the picture than just the luminance and the contrast of the foreground object and the background. To test the hypothesis that IT neurons are differentially tuned to the luminance (ΔLum_object-ground_) and contrast (ΔContr_object-ground_) of the foreground object and the background we designed a set of controlled stimuli in which we varied the luminance (Figure 3C) or contrast (Figure 3D), independently for the foreground and the background. Figure 3C (top) shows stimuli in which we parametrically varied the luminance of the foreground object and the background in opposite directions. Figure 3D shows a sequence of pictures in which we first increased the contrast of the foreground object (the four leftmost pictures) and then the contrast of the background (the four rightmost pictures). We also determined the overall picture contrast (e.g. ref.^17^), computed as the root-mean-square of the image pixels (RMS Contrast) (red curve in Figure 3C,D), which was orthogonal to the tested properties.

For each of 5 references images, we created 6 additional stimuli in which we varied the luminance of foreground and background and 6 stimuli in which we varied the contrasts of these regions (Figure 4A). As before, we presented the stimuli for 200ms, followed by 200ms with a grey screen. We carried out a linear regression to predict the MUA response to the stimuli based on ΔLum_object-ground_ and RMS Contrast for the first stimulus set and ΔContr_object-ground_ and RMS Contrast for the second stimulus set. We first normalized the responses per reference image, to discount the neurons’ tuning to shape.

**Figure 4.**
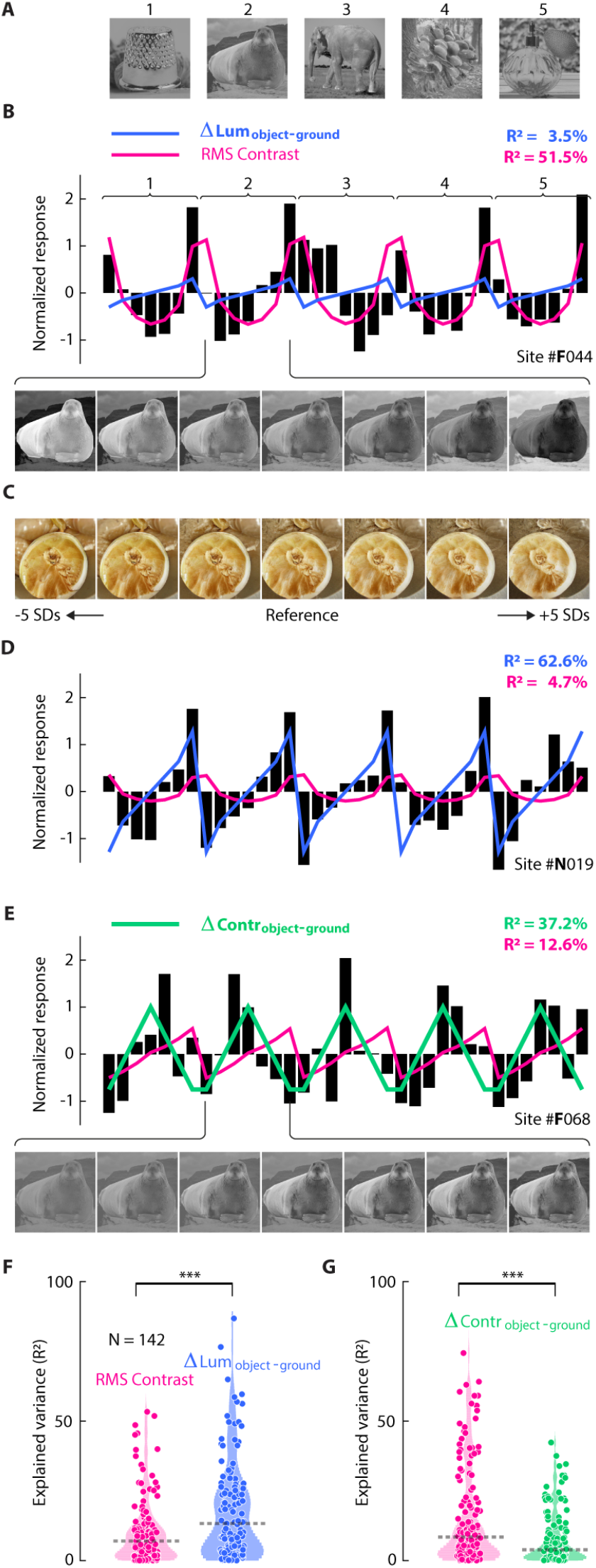
ΔLum_object-ground_ explains IT response better than contrast. We presented stimuli (for 200ms followed by 200ms grey) in which we varied the luminance and contrast of the foreground object and the background. **(A)** Five reference images were used to generate sequences with an additional six stimuli, so that total number of stimuli was of 35. **(B)** Activity elicited at an example recording site, at which the response was best predicted by RMS Contrast. **(C)** An example tuning sequence from the same site suggested that contrast has a strong influence on the response of the neurons. **(D)** Example site at which ΔLum_object-ground_ was a good predictor (same as in Figure 3B, top). **(E)** Example site at which ΔContr_object-ground_ was a good predictor (same site as in Figure 3B, bottom). **(F)** The variance of the activity across recording sites (N = 143) predicted by RMS Contrast (pink) and ΔLum_object-ground_ (blue). ΔLum_object-ground_ was a better predictor than RMS Contrast (p < 0.001, Wilcoxon signed rank test). **(G)** Same as **(F)** for the set of stimuli that were used to test RMS Contrast (pink) and ΔContr_object-ground_ (green).

Figure 4B shows the response (black bars) and ΔLum_object-ground_ (blue line) and RMS Contrast (pink line) for a representative site. The prediction by RMS Contrast was better than that of ΔLum_object-ground_ (R^2^: 51.5% v. 3.5%, p < 0.001, t-test). Interestingly, this result is reflected by the tuning dimension, which has been illustrated in Figure 4C; the rightmost images have a higher contrast both in the object and the background than the leftmost images.

Other sites had ΔLum_object-ground_ as their tuning dimension.

Figure 4D shows the results in the representative site #N019, which was also illustrated in Figure 3B. ΔLum_object-ground_ was the best predictor of the response, whereas RMS Contrast explained only little variance (R^2^: 62.6% v. 4.7%, p < 0.001, t-test). Similarly, Figure 4E shows the results for site #F068, which was illustrated in Figure 3B (top) and appeared to be tuned to ΔContr_object-ground_. The activity of the neurons was predicted by to ΔContr_object-ground_ but the difference with RMS Contrast only approached significance (R^2^: 37.2% v. 12.6%, p = 0.069, t-test).

Across the population level, ΔLum_object-ground_ explained more variance than RMS Contrast (Figure 4F; medians R^2^: 13.1% vs. 6.9%, N = 142, p < 0.001, Wilcoxon signed rank test). The result was consistent across the two monkeys (monkey F, median R^2^: 16.8% v. 7.8%, N = 89, p = 0.0066; monkey N, median R^2^: 9.0% v. 6.6%, N = 53, p = 0.0404, Wilcoxon signed rank test).

At the population level, RMS contrast explained more variance than ΔContr_object-ground_ (Figure 4G; medians R^2^: 8.4% v. 3.6%, N = 142, p < 0.001, Wilcoxon signed rank test), a finding that we replicated in both monkeys (monkey F, medians R^2^: 10.7% v. 3.8%, N = 89, p = 0.0001; monkey N, medians R^2^: 5.4% v. 3.4%, N = 53, p = 0.1429, Wilcoxon signed rank test), suggesting that it was less relevant to describe the tuning of IT neurons than RMS-contrast and ΔLum_object-ground_.

Overall, these results demonstrate that the use of GANs to determine tuning dimensions provides a data-driven method for discovering novel tuning properties of IT neurons. It was relatively straightforward to interpret the GAN-generated image sequences, which inspired image-computable metrics that we validated with controlled stimuli and explained a considerable amount of variance at the population level.

### GAN-generated image sequences predict the perceptual effects of intracortical microstimulation in area IT

We next asked whether the tuning dimensions inspired by the GAN-generated images are only related to the internal representation of neurons, or if they also influence object perception^18^. To answer this question, we used microstimulation to test the role of the neurons’ tuning dimensions in perception.

We trained the monkeys to perform an image categorization task in which they had to judge whether a sample image was more similar to one of two alternative images (Figure 5A). We presented the sample stimulus for 200ms, while the monkeys maintained gaze on a fixation point, followed by grey screen with a duration of 100ms. Then the choice display appeared and after 400ms the fixation point disappeared, cueing the monkey to make an eye movement to the image that was most similar to the sample stimulus, and the monkey was rewarded in case of correct response. To test the influence of the IT neurons on perception, we applied electrical microstimulation through one of the electrodes for 200ms, from 75 to 275ms after the onset of the sample stimulus (current: 50 µA; frequency: 300 Hz; see Methods) in 50% of the trials.

**Figure 5.**
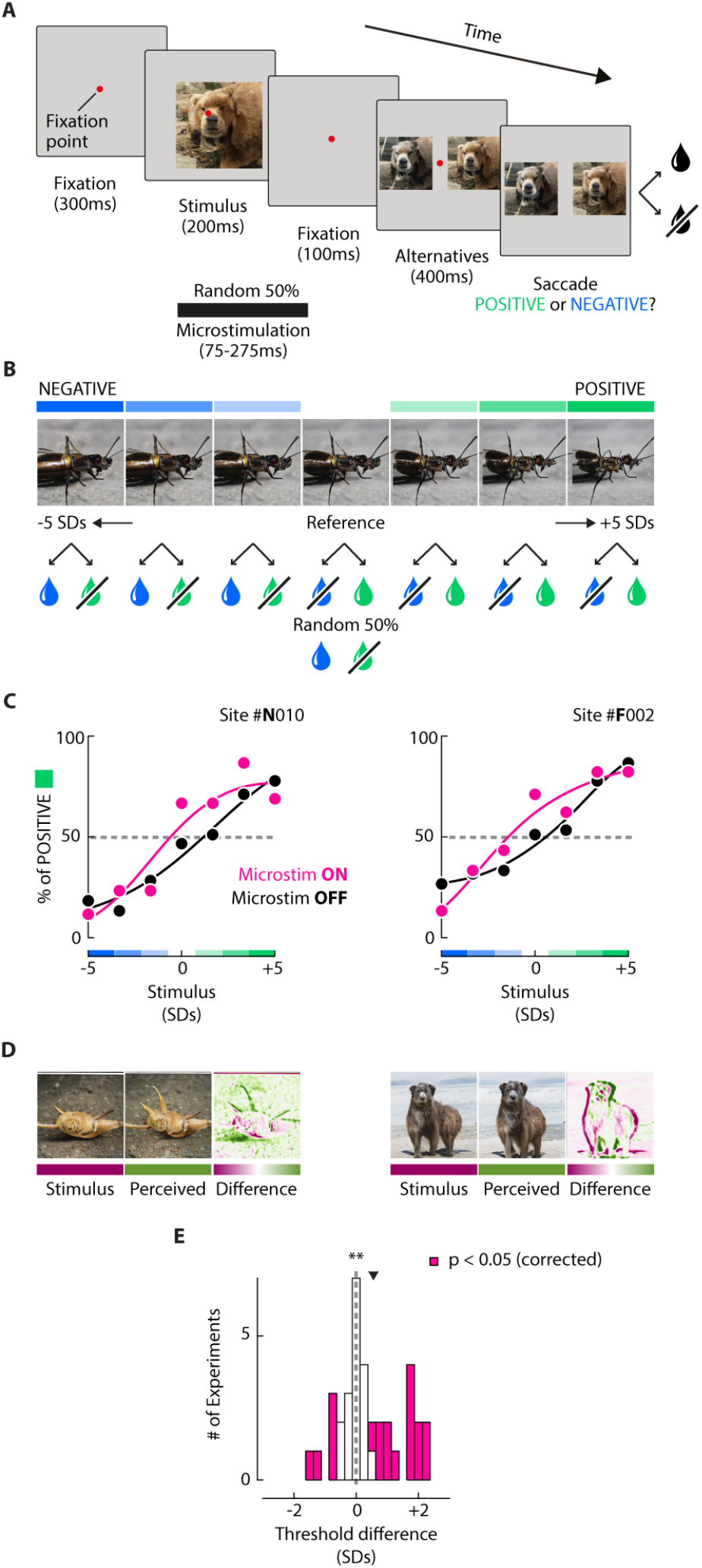
Microstimulation of IT neurons warps perception of GAN-generated images. We tested if microstimulation of IT neurons influenced the perception of the GAN-generated images. **(A)** The monkeys were trained on a delayed match to sample task. They saw a sample stimulus and were rewarded for making an eye movement to the image that was most similar to the sample. We applied microstimulation to an IT electrode on 50% of the trials from 75 to 275ms after the onset of the sample stimulus (black line at the bottom). **(B)** The sample was one of seven images generated by the GAN. The choices were the most negative (blue) and positive (green) image of the sequence and were fixed across trials. We gave a reward if the monkey chose the image that was most similar to the sample. However, if the sample was the reference images, we gave reward on a random 50% of the trials. **(C)** Choices during microstimulation of two sites, one per monkey (pink: microstimulation ON trials; black: microstimulation OFF trials). For both example sites, microstimulation biased the monkeys to choose the image that the neurons at the recording site preferred. Estimated image at which the monkey chose either image category equally often (dashed line in C), without (purple, left) and with microstimulation (middle, green). The right panel illustrates the difference between these images. **(E)** The distribution of perceptual shifts across all experiments (pink: p < 0.05, permutation test of the difference in area under the psychometric curves, 1,000 iterations, FDR corrected). We performed micro-stimulation at a total of 36 recording sites.

We performed 36 microstimulation experiments (20 in monkey N, 16 in monkey F). In each trial, the target stimulus was randomly selected from the 7 images of a GAN-generated image sequence that was based on the tuning of the recording sites at which we applied microstimulation. The two alternative images corresponded to the extremes of the sequence (Figure 5B, top) and their position was randomized across trials. We rewarded monkey for eye movement to the image that was most similar to the sample (Figure 5B). When we used the central reference image as the sample, however, we gave reward randomly on 50% of trials, irrespective of the choice. In each experiment, we used GAN-generated images derived from 4 reference images. These reference images were the same for all experiments in each monkey, but differed between the animals.

As in previous work^19–22^, we evaluated a possible effect of microstimulation on perception by fitting a psychometric function to the monkeys’ choices, separately for trials with and without microstimulation. The results of two example experiments, one for each monkey, are show in Figure 5C. In these experiments, microstimulation biased the monkey to choose the image that elicited a higher response at the recording sites. We also illustrate the images that gave rise to 50% choice of each image category, with and without microstimulation (Figure 5C). The difference between these two images visualizes the putative effect of microstimulation on the monkey’s perception (Figure 5D).

Across all the experiments, microstimulation induced a significant perceptual shift towards the image preferred by the neurons (Figure 5E; p = 0.0164, Wilcoxon signed rank test). Furthermore, 39.9% (14/36) of the experiments were associated with a shift towards the preferred image and only 13.9% (5/36) with a shift in the opposite direction (ps < 0.05, permutation test of the difference in area under the psychometric curve s, 1,000 iterations, FDR corrected), and the result was consistent in both monkeys (monkey F: 50% positive, 12.5% negative; monkey N: 30% positive, 15% negative).

In sum, we demonstrated that microstimulation warps perception in a predictable and quantifiable fashion, revealing a link between the tuning of IT neurons and their influence on conscious perception.

## Discussion

Here, we presented a novel data-driven method to infer the tuning of high-level visual neurons. This approach exploits GANs to generate image sequences that are easy to interpret and translate into novel hypotheses, in the form of image-computable metrics. We built two sets of controlled stimuli inspired by the GAN-generated images, one in which we varied the luminance difference between objects and ground (i.e., ΔLum_object-ground_) and one in which the contrast difference varied between object and ground (i.e. ΔContr_object-ground_). We found that ΔLum_object-ground_ predicted a significant amount of variance at the population level, while ΔContr_object-ground_ explained the activity only of a few recording sites. In addition, we have shown that the GAN-generated image sequences can predict the effect on perception indunced by electrical microstimulation of IT neurons.

The performance of ΔLum_object-ground_ was not totally unexpected: two previous studies reported that a few shape-selective^23^ and face-selective^24^ neurons preferred a specific contrast polarity, and its reverse would reduce their response, although further studies did not find a consistent role of contrast inversion in IT^25,26^. However, these previous studies provided only a limited understanding, based on a binary definition of contrast polarity and not a systematic investigation of the linear relationship between this property and the response of IT neurons, independently of shape-selectivity, and using natural stimuli. For this reason, we considered that testing this property would potentially lead to the discovery of a novel form of tuning in IT neurons. Indeed, this proved a surprising result because luminance is one of the most basic visual properties, and as such, we expected high-level visual neurons in IT to be invariant to it. Of course, this form of tuning relies on the association of luminance with figure-ground organization, which is indeed a high-level visual property. Still, this finding raises the question of what cortical pathway leads to this form of tuning, as luminance-invariance emerges as early as V1 in the visual cortex (e.g. ref.^27^). These IT neurons might infer object-ground organization by reading out the differential activity of, e.g. thalamic visual neurons tuned to luminance, or they might inherit their selectivity from upstream regions with direct connections to luminance selective neurons.

Another interesting result is that the GAN-generated image sequences varied along multiple dimensions. We confirmed this finding using controlled stimuli, because different image properties independently explained variance in the activity of neurons at several recording sites (e.g. in Figure 4E), in line with findings on the tuning of IT neurons to multiple properties^24,28–34^.

Our approach can be used to test other hypotheses about the dimensionality of cortical representations, and the entanglement of tuning dimensions^35–37^. Indeed, it was exactly by looking at this that Higgins et al.^35^ developed a similar approach to ours, to retrieve the tuning dimensions of face-selective neurons in IT, exploiting a GAN trained with the objective to learn a disentangled latent space, whose dimensions varied along interpretable features such as age and gender. However, the present results go beyond their work by validating the feature to which the IT neurons are tuned to with controlled stimuli and by testing the influence of the tuning of IT neurons on perception with microstimulation. Indeed, we found that the GAN-generated image sequences allowed us to influence perception in a predictable and quantifiable fashion with microstimulation, determining a significant shift in perception for ∼40% of the electrodes. This percentage reflects the uncertainties on the local effects of microstimulation, that might impact a slightly different population than the one being recorded (e.g. because of nearby axons being stimulated as well)^18^, the different detection thresholds in different sites, and the fact that despite the use of chronic implants, the overlap between the population recorded on the first day and the population microstimulated weeks later might be too low for certain sites. However, our results are in line with previous studies that used a similar paradigm in many different regions, all reporting significant effects in ∼30-60% of the microstimulation experiments^19,20,22^.

This form of AI-driven microstimulation may also prove beneficial for future visual cortical prostheses to enrich the perception of the user by directly influencing how an object is perceived. Our work demonstrated that GAN-generated sequences are useful for predicting the tuning of high-level visual neurons and even controlling conscious perception.

We will now address some the limitations of the method. Firstly, it is only possible to determine the tuning of the neurons to features that are captured by the latent space of the GAN, which depends on the set of stimuli that are used to train it. We expect that the importance of this limitation will decrease over time, because the quality of the images that can be generated by ANNs is increasing steadily, and their latent spaces are becoming broader. Importantly, the proposed method is fully compatible with these newer models.

Second, the GAN-generated image sequences that were based on the tuning of individual recording sites varied across multiple dimensions, so that additional and better controlled stimuli are necessary to confirm the tuning of the neurons to specific features. The necessity of the validation of the tuning is shared with other methods, including MEIs^5,6^.

Third, we recorded multi-unit activity, and we did not isolate individual neurons. Hence, the GAN-generated images sequences gave insight into the average tuning of multiple neurons at each of the recording sites. We note, however, that the method can also be used to determine the tuning of single neurons. Recording MUA might, however, be advantageous if the goal is to predict the influence of microstimulation, because it may provide a better approximation of the tuning of the set of neurons that are influenced by microstimulation.

In the present study, we focused on the early responses elicited by the onset of the stimuli: we aimed at providing a proof of concept that the method could be used to generate interpretable image sequences, potentially leading to novel hypothesis on the tuning of high-level visual neurons. However, future studies could examine the relation between the latent GAN space and the later response phase of IT neurons. Furthermore, future research might use the present method to study the topographic organization of the tuning dimensions, an approach that could also be used in combination of imaging techniques, including 2-photon imaging and fMRI. In addition, the method might be extended to movies and reveal insight into the tuning of units across successive frames, like for example the tuning to different types of motion, including biological motion.

In conclusion, we illustrated a data-driven method to study the tuning of IT neurons that can be used to find sensitivity of the neurons to features that might not have been anticipated by previous work.

## Materials and Methods

### Experimental model and subject details

All procedures complied with the NIH Guide for Care and Use of Laboratory Animals and were approved by the “Centrale Commissie Dierproeven” of the Netherlands. Two macaque monkeys (males, 7 and 6 years old) participated in the experiments. They were socially housed in stable pairs in a specialized primate facility with natural daylight, controlled humidity and temperature. The home-cage was a large floor-to-ceiling cage which allowed natural climbing and swinging behaviour. The cage had a solid floor, covered with sawdust and was enriched with toys and foraging items. Their diet consisted of monkey chow, supplemented with fresh fruit. Their access to fluid was controlled, according to a carefully designed regime for fluid uptake. During weekdays, the animals received water or diluted fruit juice in the experimental set-up upon completed trials. We ensured that the animals drank sufficient fluid in the set-up and supplemented the animals with extra fluid after the recording session if they did not drink enough. During the weekend, they received a full bottle of water (700-940 ml per day) in the home cage. The animals were regularly checked by veterinary staff and animal caretakers and their weight and general appearance were recorded daily in an electronic logbook during fluid-control periods.

### Surgeries

Each animal received three cranial implants during separate surgical procedures. The implants were customized, in-house-designed, 3D-printed titanium components^38^. In the first surgery we implanted a head post for head fixation, ensuring that the eye tracker could capture the eye position. After several months, to allow bone growth and a firm attachment to the skull, we implanted a baseplate in a second surgery. The base plate was implanted under the skin and served as base for the 1024 channel connector. The third surgery followed after several months. We implanted 16 Utah electrode arrays (Blackrock Microsystems), attached with wire bundles to a customized 3D-printed titanium pedestal with a 1024 channel connector that was secured with screws to the baseplate. For the present study we only used the recordings from the IT arrays.

### Electrophysiology

We recorded neuronal activity from 256 recording sites in IT in monkey N (4 Utah arrays) and 320 in monkey F (5 Utah arrays). The amplified signal was sampled at 30kHz using a Blackrock Microsystems system. Neural signals were referenced to a subdural electrode and amplified using Blackrock Microsystems Cereplex-M headstage amplifiers. We measured the envelope of multi-unit activity by band-pass filtering the signal offline (2nd order Butterworth filter, 500 Hz-5 KHz, filtfilt.m in MATLAB) to isolate high-frequency (spiking) activity. This signal was rectified (negative becomes positive) and low-pass filtered (corner frequency = 200 Hz) to produce the envelope of the high-frequency activity, which we refer to as MUA^39^. The MUA signal was down-sampled to 1kHz and stored for further analysis. The MUA signal reflects the population spiking of neurons within 100-150 µm of the electrode and the population responses are very similar to those obtained by pooling across single units^39,40^.

### MUA pre-processing

We averaged the MUA signal for each recording site and each trial in the time-window 75-175ms after the stimulus onset and then we normalized the data (as in ref.^10^). For each day of recording, we computed the mean and standard deviation of the activity elicited by the test images (GAN experiment) or by all the stimuli (controlled experiment). We called this sub-group of trials ‘*test pool’*. The normalized MUA for each trial *t* and recording site *n* was then calculated as follows:

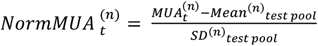

To estimate the reliability of the MUA signal at each recording site, we computed the mean reliability^10^, defined as the average of the pairwise correlations between each combination of repetitions to the same images. We selected recording sites with a mean reliability higher than 0.3 for the GAN experiment (following: ref.^10^), and 0.75 for the experiment with controlled stimuli. During the microstimulation experiments, we connected a stimulation device (Cerestim - Blackrock microsystems) to one bank of IT electrodes (32 recording sites) and selected one site for stimulation per experiment. On microstimulation trials, we applied 60 bipolar pulses of 50 µA at a frequency of 300Hz (pulse-width = 170 µS, interphase-delay = 60 µS). A subdural electrode was used for the current return.

### Stimulus presentation

We used a CRT monitor with refresh rate of 60Hz and resolution of 1024×768 pixels, which was viewed from a distance of 58cm (26.9 pixels per degree). Stimuli were generated in Matlab using the COGENT Graphics toolbox (developed by John Romaya at the LON at the Wellcome Department of Imaging Neuroscience) and custom control software^41^. To ensure that all RFs would fall inside the image, we shifted the stimuli 100 pixels downward and 100 pixels to the right relative to the centre of gaze. In all experiments, we pseudorandomized the order of presentation of images. In total, we recorded 19,675 complete trials from monkey N (GAN experiment: 2,000 trials; Controlled experiment: 875 trials; 20 microstimulation experiments: 16,800 trials) and 16,315 trials in monkey F (GAN experiment: 2,000 trials; Controlled experiment: 875 trials; 16 microstimulation experiments: 13,440 trials).

### Passive fixation task

The first two experiments (GAN and controlled experiment) used a passive fixation task. At the start of the trial, the monkey directed its gaze to a red fixation point with a diameter of 0.3 deg on a grey background (luminance 15.7 cd/m^2^). We presented the first image once the monkey had maintained fixation for 300ms in a fixation window with a diameter of 1dva. To receive a reward, the animal had to maintain its gaze in this window during the sequence of four images, which were each presented for 200ms, followed by 200ms with a grey screen. We logged images as completed when the monkey maintained fixation during their presentation and during the following epoch of 200ms with a grey screen. If trials were aborted because the monkey broke fixation, completed images were not shown again, and the non-presented images of the aborted sequence were presented on the next trial.

### Delayed match to sample task

The microstimulation experiments required a perceptual similarity judgment. At the start of the trial, the monkey directed its gaze to a red fixation point with a diameter of 0.3 dva on a grey background (luminance 15.7 cd/m^2^). We presented the first image, which was the sample stimulus, once the monkey had maintained fixation for 300ms in a fixation window with a diameter of 1dva. The animal had to maintain its gaze in the fixation window during the 200ms that the sample stimulus was visible, followed by 100ms of a grey screen, and the choice display with two pictures, each on one side of the screen. The side of the two pictures was (pseudo-) randomized across trials. After another 400ms, the fixation dot disappeared, cueing the monkey to make a saccade to the choice image that was most similar to the sample. In 50% of the trials, we delivered microstimulation to a specific site from 75 to 275ms after the onset of the sample stimulus. Reward was delivered in case of correct response but at random for the reference images, at the centre of the image sequence. We repeated the trials that were aborted because the monkey broke fixation.GAN experiment: network.

We replicated the same methods as in ref.^10^ In brief, we employed StyleGAN-XL which has been successfully trained on ImageNet^42^ to generate high-resolution images of a thousand different categories, resulting in a complex and diverse stimulus dataset.

This GAN maps the input latent vector “z” (z-latent) to an intermediate latent vector “w” (w-latent) to favor feauture disentanglement. The original z-latent space is restricted to follow the data distribution that it is trained on and it is conditional to a specific class present in ImageNet. However, the w-latent space overcomes this limitation and it is also not conditioned to single ImageNet classes.

### GAN experiment: stimuli

We synthesized images from the 200 classes from Tiny ImageNet (a subset of the thousand classes from ImageNet)^43^ so that each class was represented by twenty training set stimuli and one test set stimulus.

First, a 64-dimensional vector was sampled from a standard Gaussian and concatenated with the 64-dimensional embedded representation of the class category, resulting in 128-dimensional z-latents that were utilized to synthesize RGB images with a resolution of 512 × 512 pixels. For the training set, z-latents were randomly sampled and mapped to w-latents that were truncated at 0.7 to support image quality as well as diversity. The average w-latent of each category was utilized for the test set (see ref.^10^). In total, the training and test set consisted of 4000 stimuli, that were each presented once and 200 stimuli that were presented 20 times, respectively. Finally, we performed a multiple linear regression to learn a linear mapping between the w-latent of the training set and the MUA responses evoked by the training images in IT (see main text).

Latent vectors are available online at: <u>10.6084/m9.figshare.25637856</u>. Code to generate the stimuli is available online at: https://github.com/neuralcodinglab/brain2gan. The MUA responses will be available online upon paper acceptance.

### Controlled experiment: stimuli

We selected five pictures of objects and five compatible natural scenes from copyright-free images on *Pixabay*. We cropped the objects and placed them in the natural background, ensuring that the foreground objects and backgrounds had the same number of pixels. For the stimuli in which we varied ΔLum_object-ground_, we changed the luminance of the object and the luminance in opposite directions. In the 7 steps, we summed the following values to all the object pixels, transformed into percentages, where 0 is the mean of the image portion: -60, -30, -15, 0, 15, 30, 60; and the following ones to all the ground pixels: 60, 30, 15, 0, -15, -30, -60. To vary ΔContr_object-ground_ we changed the contrast of the foreground object and background independently. In the 7 steps, we multiplied all the object pixels by the following percentage: 50, 75, 100, 125, 125, 125, 125; and changed all the ground pixels by the following percentage: 50, 50, 50, 50, 75, 100, 125. Finally, RMS contrast was computed as follows:

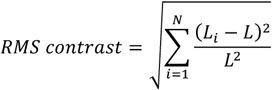

Where *N* is the number of pixels in the stimulus. *L* is the mean luminance of the pixels and *L*_*i*_ is the luminance of the *i*-th pixel.

We presented each stimulus 50 times and averaged the MUA across repetitions. Some of the neurons responded more strongly to some of the five pictures that to others. To account for this shape selectivity, we z-scored the MUA responses within each image sequence (one sequence per reference image) and performed one multiple linear regression for each recording site, separately for the experiment in which we varied ΔLum_object-ground_ and ΔContr_object-ground_, to test the influence of the predictors on the magnitude of the MUA responses in IT neurons.

The stimuli and MUA responses will be available upon paper acceptance.

### Microstimulation experiments: GAN-generated image sequences

We created the GAN-generated image sequences as described in the above. We selected 12 arbitrary reference images for each monkey, including animal faces, entire animals, tools, insects, food and both indoor and outdoor scenes. Then we produced the image sequence for 75 sites in monkey F, and 161 sites in monkey N for each of the 12 reference images. For the microstimulation experiments, we further selected image sequences based on 4 reference images from different semantic classes (these classes also differed between the monkeys). In our selection of recording sites for microstimulation, we chose recording sites with a high reliability (in the GAN experiment): we computed the reliability as the split-half correlation across trials, and select sites with a correlation higher than 0.3. We did not select sites that were close to each other (<800 µm). We tended to select channels within the same 32-channel bank on successive days, to avoid unnecessary cable disconnections/connections which can damage the connectors.

### Microstimulation experiments: analysis of behaviour

For each experiment, we determined the percentage of responses of the two classes for every stimulus. We expected microstimulation to induce a higher percentage of responses to the positive image, thus we computed the difference in area under the psychometric curve (AUC) of each condition^44^: if microstimulation was effective, the area under the curve computed using trials with microstimulation ON would have been larger compared to that computed using trials without microstimulation. We computed the statistical significance of the difference in AUC between conditions using a permutation test. In brief, for the same stimulus, we randomly scrambled the trials in the two conditions 1,000 times, and then we computed the difference in AUC for each scrambled configuration. The p-value was computed as the percentage of scrambled AUC differences higher than the true AUC difference. Finally, we applied the family-discovery-rate algorithm to correct for multiple correction across experiments^45^.

To quantify the change in perception, we fitted a psychometric function using a model-free non parametric estimation^46^, for each experiment and condition separately, and then computed the image giving rise to an equal number of reports to the two classes, with and without microstimulation (Figure 5D). This allowed us also to test the effect at the population level (Figure 5E). Notably, we found a big overlap in significant experiments when computing selectivity using a more traditional statistical test based on the shift between psychometric curves (as in ref.^47^), or when excluding experiments based on the behavioural performance (not shown). We report in the main text the values for the AUC-difference, as they reflect the actual recorded data and the tests were more conservative.

## Acknowledgements

We thank Kor Brandsma, Anneke Ditewig, Taijsha van Rees and Lex Beekman for biotechnical support; Maurice Heemskerk and Mike Vink for technical assistance.

The work was supported by NWO grants (Crossover grant 17619 “IN-TENSE” and “DBI2”, a Gravitation program of the Dutch Ministry of Science), the European Union Horizon 2020 Framework Program under specific grant agreement 945539 “Human Brain Project”, grant agreement 899287 “NeuraViper,” an ERC grant (101052963 “NUMEROUS”) to PRR and by NWO grants (OCENW.XS22.2.097 and VI.Veni.222.217) to PP.

## Author contributions

PP and PRR conceived of the study. PP and DDL performed the experiments. PP analysed the data. PP and PRR obtained funding. PP and PRR wrote the paper with input from DDL.

## Notes

### Competing Interest Statement

The authors have declared no competing interest.

